# Real-time functional connectivity-based neurofeedback of amygdala-frontal pathways reduces anxiety

**DOI:** 10.1101/308924

**Authors:** Zhiying Zhao, Shuxia Yao, Keshuang Li, Cornelia Sindermann, Feng Zhou, Weihua Zhao, Jianfu Li, Michael Lührs, Rainer Goebel, Keith M. Kendrick, Benjamin Becker

## Abstract

Deficient emotion regulation and exaggerated anxiety represent a major transdiagnostic psychopathological marker. On the neural level these deficits have been closely linked to impaired, yet treatment-sensitive, prefrontal regulatory control over the amygdala. Gaining direct control over these pathways could therefore provide an innovative and promising strategy to regulate exaggerated anxiety. To this end the current proof-of-concept study evaluated the feasibility, functional relevance and maintenance of a novel connectivity-informed real-time fMRI neurofeedback training. In a randomized within-subject sham-controlled design high anxious subjects (n = 26) underwent real-time fMRI-guided training to enhance connectivity between the ventrolateral prefrontal cortex (vlPFC) and the amygdala (target pathway) during threat exposure. Maintenance of regulatory control was assessed after three days and in the absence of feedback. Training-induced changes in functional connectivity of the target pathway and anxiety ratings served as primary outcomes. Training of the target, yet not the sham-control, pathway significantly increased amygdala-vlPFC connectivity and decreased subjective anxiety levels. On the individual level stronger connectivity increases were significantly associated with anxiety reduction. At follow-up, volitional control over the target pathway and decreased anxiety level were maintained in the absence of feedback. The present results demonstrate for the first time that successful self-regulation of amygdala-prefrontal top-down regulatory circuits may represent a novel strategy to control anxiety. As such, the present findings underscore both the critical contribution of amygdala-prefrontal circuits to emotion regulation and the therapeutic potential of connectivity-informed real-time neurofeedback.

## Introduction

Successful regulation of negative affect is crucial for mental health and well-being (1, 2). Deficient emotion regulation (ER) and exaggerated anxiety represent transdiagnostic markers across major psychiatric disorders, including the most prevalent Axis I disorders such as anxiety and addiction as well as Axis II disorders ((3–5), see also recent meta-analysis (6)).

The functional interplay between and clinical relevance of ER and anxiety mirrors across different levels of observation. In healthy subjects, ER capability prospectively predicts anxiety levels for periods up to five years (7–9). The clinical relevance is further emphasized by randomized trials evaluating the efficacy of cognitive-behavioral therapy (CBT) indicating that improved ER predicts symptom reduction in anxiety disorders (10–12). On the neural level efficient regulation of threat and anxiety is neurally underpinned by top-down governance of the amygdala, which is critically engaged in threat responsivity (13), via prefrontal regulatory regions (14, 15). Within these regulatory circuits the ventrolateral (vlPFC) and dorsomedial (dmPFC) prefrontal cortex are considered to specifically support explicit/volitional control of threat via downregulation of the amygdala (5, 15, 16). Deficits in this top-down regulatory mechanism have been identified across major psychiatric disorders (17), with disorders characterized by exaggerated anxiety exhibiting decreased recruitment the prefrontal cortex and concomitantly exaggerated amygdala activity in the context of attenuated functional interplay between these regions (17–19). The therapeutic relevance of these pathways is further emphasized by studies reporting that anxiety reduction following behavioral and pharmacological interventions is accompanied by normalization of deficient amygdala-prefrontal coupling (20–22).

Despite the important contribution of neuroimaging research to identifying altered amygdala-prefrontal interaction and its normalization as a potential pathological and treatment-sensitive neural marker for neuropsychiatric disorders characterized by emotional dysregulations, it has yet to directly have a therapeutic impact (23). Given that the currently available therapeutic interventions for anxiety reduction are generally characterized by moderate response rates and potential negative side effects (24–26), innovative treatments that directly target the identified brain markers are needed (27). Within this context, the emergence of real-time fMRI neurofeedback (rt-fMRI NF) training approaches that allow subjects to gain volitional control over regional brain activity have been considered as a putatively promising strategy (28–30). Importantly, previous studies have confirmed this potential of rt-fMRI NF by demonstrating that training success in terms of control over regional activity can be maintained beyond the training session (31–34), and that training-induced neural activity changes can modulate emotional experience in healthy subjects (31) and patients with major depression (32–35).

Initial studies have begun to evaluate the therapeutic potential of rt-fMRI NF in clinical populations and demonstrated that up-regulating activity in primary emotion processing regions such as the insula and amygdala can successfully decreased symptoms in patients with major depression (32–34). Given the critical role of the amygdala in anxiety and consistently observed hyper-responsivity in this region in anxiety-related disorders (18, 36, 37) previous rt-fMRI NF studies trained subjects to down-regulate neural activity in this region and demonstrated that this strategy has the potential to enhance ER and attenuate anxious arousal (38–41). In line with current neuro-circuitry models of ER, successful down-regulation of the amygdala was accompanied by increased functional connectivity between the amygdala and prefrontal regulatory regions in both, healthy subjects (39, 42) as well as patient populations with exaggerated anxiety (40, 43).

Summarizing, the current literature suggests that (a) successful ER relies on top-down regulation of the amygdala via prefrontal regions and that (b) rt-fMRI NF-assisted modulation of these regions has the potential to modulate ER and anxious arousal. In the context of recent circuit level models of ER (for circuit-level deficits in psychiatric disorders see (44)), the present randomized sham-controlled within-subject proof-of-concept study aimed at evaluating whether (1) rt-fMRI NF has the potential to directly allow regulatory control of the strengths of functional connectivity in the amygdala-prefrontal regulatory pathways, (2) successful regulatory control decreases levels of anxiety in individuals with high anxiety, and (3) volitional control can be maintained in the absence of feedback and over a period of three days.

## Methods and Materials

### Participants

To increase the clinical relevance of the present proof-of-concept study while controlling for potential confounding factors in clinical populations, including co-morbidity or medication, healthy subjects with high anxiety (trait anxiety scores > 40, assessed by STAI (45)) were recruited. Given that the main aim was to evaluate the feasibility and functional relevance of connectivity-based rt-fMRI NF training potential confounding effects of menstrual cycle-related variation in ER (46), as well as sex-differences in emotion regulation (47) and associated connectivity in the target pathway (48) were controlled for by focusing on a male sample. Detailed eligibility criteria and sample characteristics are provided in the **Supplementary Materials**. All subjects provided written informed consent. The study had full ethical approval by the local ethics committee, adhered to the latest reversion of the Declaration of Helsinki and protocols were pre-registered (NCT02692196, http://clinicaltrials.gov/show/NCT02692196).

### Protocols and procedures

Participants were scheduled for four MRI sessions: rt-fMRI NF training of the amygdala-vlPFC target pathway (EXP) plus transfer/maintenance assessment after two days (M-EXP) and rt-fMRI NF sham (SHC) training plus transfer/maintenance assessment after two days (M-SHC). During the training sessions feedback was provided but not during the transfer/maintenance sessions. Training sessions used identical procedures including four subsequent NF training runs and during the sham session participants received connectivity feedback from a pathway connecting regions not engaged in ER (bilateral motor cortices, M1 (15)). The SHC served to control for unspecific effects of training, and together with the within-subject design allowed a thorough control of potential confounders. To control for carry-over effects the order of training sessions was counterbalanced. For randomized allocation of the order of trainings a random number generator was used. Training sessions were separated by an interval of 2-3 weeks and subjects were informed that they had to discover new strategies each time. Both training sessions were preceded by fMRI-localizer paradigms to determine the pathways used for feedback during EXP and SHC (see **Localizer-paradigms**). Two days after each training session, participants underwent two transfer runs (M-EXP/M-SHC) during which they were required to perform regulation with the same strategy they had learned during the preceding training but without feedback being provided (protocols see **Figure 1**). To evaluate the functional relevance of training success, anxiety levels assessed before and after each session and served as primary behavioral outcome. To control confounding effects of pre-training mood and anxiety states these were assessed additionally immediately before each training and maintenance session.

### Localizer-paradigms

Pathway-specific localizer paradigms were employed to localize the target emotion regulation nodes (EXP, right amygdala and right vlPFC, **emotion localizer**) and the bilateral motor cortices (SHC, bilateral M1, **motor localizer**). These regions-of-interest (ROIs) for the feedback pathways were determined using a combined structure-function approach. Thus, T1-weighted brain-structural images of each subject were overlaid with the real-time localizer activity in native space. Subsequently, thresholded peak activity regions within the target structures were manually delineated (standardized ROI size) and used for the subsequent training. To further control for movement effects on functional connectivity (49) and physiological artifacts such as respiration and noise from cardiac activity, a third ROI was placed in a right postcentral white matter tract (**Supplementary Materials**).

### Neurofeedback training protocols

During the neurofeedback training (NFT) strong negative (threatening) stimuli were displayed with real-time neurofeedback displayed by thermometer bars on both sides (**Figure 1**, stimuli details see **Supplementary Material**). Each of the four NFT runs included four blocks of threatening pictures (six pictures per block, inter-block interval 30 seconds fixation, pictures presented for 5 seconds, size gradually increased from half to full size stepped by the TR to increase threat). Rt-fMRI connectivity (FC) between the ROIs was calculated as a partial correlation between the time series from the two pathway ROIs (amygdala-vlPFC or bilateral M1) while including the time series from the white matter ROI as a covariate (details see (50)). The FC thermometer was updated in real-time (logged to the TR = 1.5 seconds).

Participants were informed that the purpose of the training was to enhance their emotion regulation abilities to improve coping with negative emotional events in daily life and reduce stress. Participants were instructed to learn to control the threatening feelings evoked by the pictures while breathing normally. To increase their regulation ability, the neural emotion regulation success would be presented to them (thermometer bars corresponding to better success) and they should aim to develop a strategy to increase the bars. As a specific strategy is not necessary for successful learning in NFT (51, 52), no strategies for emotion regulation were introduced to the participants except that they should not try to increase the thermometers by physical means such as low breathing or head/body motion. Participants were encouraged to discovery an efficient strategy to regulate the bars and control threatening feelings. Once they discovered an efficient strategy to increase the feedback bars they were asked to continue using it during the subsequent training and transfer sessions. Finally, subjects were informed that the feedback would be computed in real-time but displayed with approximately 10 seconds delay (see also **Evaluation of training success on the neural level**)

**Figure 1.**
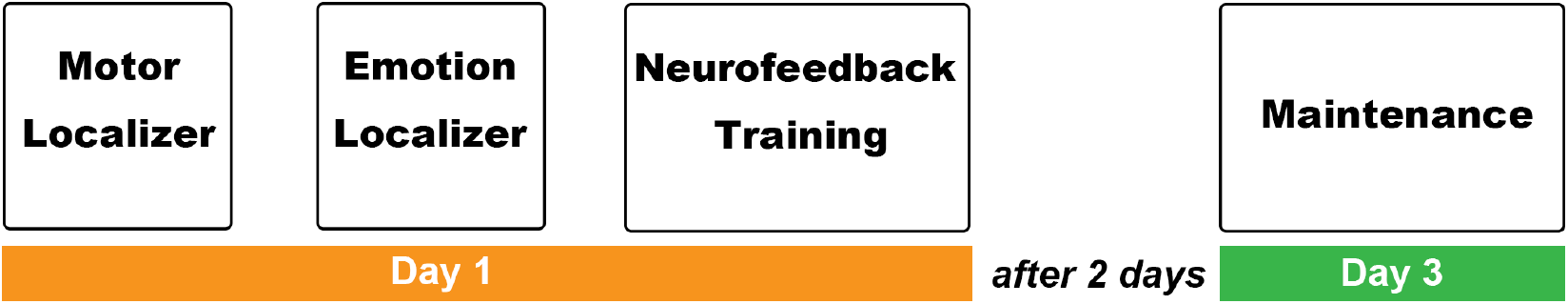
Experiment procedures for both training sessions (EXP/SHC). Training sessions for each participant were separated by a 2~3 week interval.

### Primary behavioral outcome and control variables

The primary behavioral outcome to evaluate the functional relevance of the target pathway training was determined as training-induced changes in subjective anxiety levels as assessed by visual analog scales (VAS, anxiety levels from 0 to 100) administered before and after each training session. To further control for confounding effects of pre-training differences in mood and state anxiety between the sessions, corresponding indices were assessed by means of the PANAS (the Positive and Negative Affect Schedule (53)) and SAI (State Anxiety Inventory (45)) administered before each training and maintenance session. Behavioral measures were assessed outside of the scanner by an experimenter blinded to the training condition (EXP, SHC).

### MRI Data acquisition, online preprocessing and connectivity neurofeedback

Data was acquired at 3 Tesla using evaluated sequences (**Supplementary Materials**). Online data preprocessing and real-time feedback were computed using Turbo-BrainVoyager v3.2 (Brain Innovation, Maastricht, The Netherlands). To increase the signal-to-noise ratio of the functional data during online processing real-time preprocessing was applied including motion correction and spatial smoothing with a 4mm Full Width at Half Maximum (FWHM) Gaussian kernel and temporal drift removal applied as confound predictor to the GLM. Based on findings from a previous study (54) a sliding window approach with a length of 7.5 seconds (5 volumes per window, TR 1.5 seconds) was chosen to compute the real-time connectivity feedback. Feedback was thus provided as a partial correlation coefficient between the two ROI time series segemented in consecutive windows while controlling for the nuisance signal from the third ROI (**Supplementary Materials**).

### Offline preprocessing and analyses

Preprocessing for offline analysis was conducted using standard procedures in SPM12 (Statistical Parametric Mapping, http://www.fil.ion.ucl.ac.uk/spm/). To evaluate BOLD level changes during the localizers on the group level first level General Linear Models were built. To increase the sensitivity of the offline connectivity analysis individual ROIs from the training sessions were exported (**Supplementary Materials, Table S2, Figures S1-S2**).

### Primary neural outcome and evaluation of training success

To evaluate whether NFT increased functional connectivity in the emotion regulation circuit, task-based functional connectivity was employed using a generalized form of context-dependent psychophysiological interaction (gPPI) (55) implemented as whole-brain connectivity models with the individualized amygdala (**Table S2**) or M1 ROIs used during training as seeds. The gPPI models were built on the first-level by adding the time-series from the seed region as a new regressor into the GLM design matrix. A previous study showed that a 12-second time window had comparable sensitivity in detecting task-relevant connectivity changes as a longer time window (26s) for a finger tapping task (56). Since valid online connectivity feedback started at ~12.5 seconds (5 seconds delay from the BOLD response, and 7.5 seconds length of the slide time window) after the start of a regulation block, analysis of NFT task-based functional connectivity focused on the second half of the regulation blocks (last 15 seconds of every training block - additional analyses using the entire block lengths fully confirmed the findings, **Supplementary Material**).

To evaluate training-induced changes in the target pathways connectivity strengths per run were extracted from the corresponding region (vlPFC/right M1) using beta-estimate maps generated in gPPI analysis. Estimates were further analyzed using SPSS by means of repeated measure ANOVAs and post-hoc tests controlled for multiple comparisons with Bonferroni-correction. In line with our previous study we additionally evaluated training success by comparing differences in connectivity strengths between the first two and the last two training runs (31).

## Results

### Data quality assessment protocols

Three subjects did not display above-threshold activity in the vlPFC during the emotional localizer and their data was thus excluded from all analysis, resulting in n = 23 for the final analyses. Five runs showed > 2.5mm or > 2.5° head motion, data from these runs was consequently excluded. Head motion (mean frame-wise displacement (57)) did not differ between the experimental and the sham training (EXP vs. SHC, t_22_ = 0.58, p = 0.586).

### Mood states and anxiety

Mood and anxiety data for one participant was lost (pre-training assessment, EXP). Examination of the pre-training data from the remaining participants confirmed the recruitment of subjects with high anxiety (reflected in high state anxiety scores) and did not reveal differences in pre-training anxiety and mood between training sessions (**Table1**, paired t-test, p > 0.38).

**Table 1.** Mood states before both training sessions. Mean scores of positive and negative mood and anxiety levels and their SDs (in brackets) assessed by questionnaires. Abbreviations: PANAS-P, the Positive and Negative Affect Schedule – positive; PANAS-N, the positive and negative affect schedule – negative; SAI, State-Trait Anxiety Inventory – state anxiety.

|  | Before EXP training<br>(N = 22) | Before SHC training<br>(N = 23) |
| --- | --- | --- |
| PANAS-P | 24.95 (4.84) | 22.22 (5.38) |
| PANAS-N | 14.86 (4.02) | 14.48 (4.23) |
| SAS | 38.05 (7.06) | 36.57 (7.06) |

### BOLD response during the localizer and NFT

Group-level analysis of the localizer tasks revealed that the emotional localizer reliably activated the emotional brain networks, including the amygdala and vlPFC (SPM one-sample t-test, whole-brain, False Discovery Rate (FDR) (58), corrected p < 0.01, **Figure S1**). As expected, the motor localizer reliably activated the motor networks including bilateral M1 (**Figure S2**). Importantly, emotion regulation (Regulation - Baseline) during both EXP and SHC training induced a similar activity pattern as in the emotion localizer tasks in the ER brain networks, including dmPFC, vlPFC, amygdala, insula, and parietal regions (59). Furthermore, activity patterns overlapped with the amygdala and vlPFC ROIs derived from the emotion localizer yet not the M1 sham ROIs (**Figure 2**), further validating the M1 regions as a suitable sham-control.

**Figure 2.**
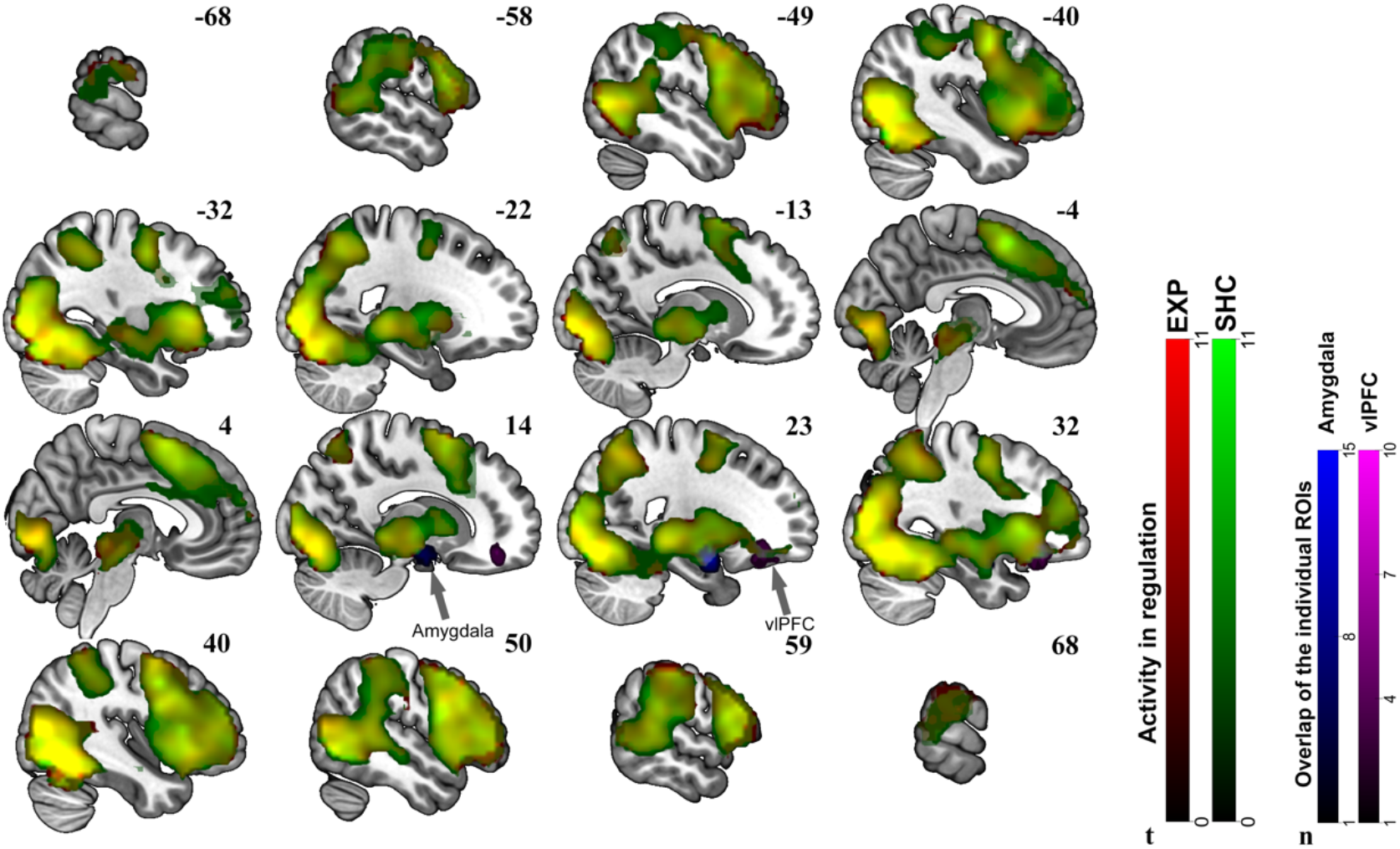
BOLD responses during NFT tasks in EXP (in red) and SHC (in green) sessions and the ROIs of EXP session. Brain activity was thresholded with p < 0.01, FDR correction. T values from SPM are indicated by corresponding color bars. The overlap between individual sphere ROIS built in offline analysis are displayed for amygdala (in blue) and vlPFC (in violet).

### Evaluation of training success on the neural level

Examining changes in functional connectivity between right amygdala and vlPFC across the four NFT runs with one-way ANOVAs (repeated measures) revealed a significant difference between them (F_3, 66_ = 3.33, p = 0.025). Importantly, this pathway did not show significant changes across the training runs with sham feedback (F_3, 54_ = 1.01, p = 0.393) (**Figure 3**). Post hoc tests for the EXP training revealed that connectivity in the target pathway did not show changes between the early training runs (Run1 vs Run2, t_22_ = 1.22, p = 0.240), but increased significantly after the second NFT run (Run3 vs. Run2, t_22_ = 2.95, p = 0.007, Cohen’s *d* = 0.78; Run4 vs. Run2, t_22_ = 3.83, p = 0.001, Cohen’s *d* = 1.04, paired t-tests, both significant after Bonferroni corrected p = 0.05/6). Again, concordant analysis of the sham training data did not yield significant changes in this pathway (t_21_ = −0.52, p = 0.607) (**Figure 3**). In line with our previous study (31) we additionally confirmed training success using a more robust estimation which compared early versus late runs in the target pathway (Run3 + Run4 > Run1 + Run2, t_22_ = 2.81, p = 0.010, Cohen’s *d* = 0.79, paired t-test). In an additional control analysis, training success was confirmed using the data from the entire blocks (see **Supplementary materials**).

**Figure 3.**
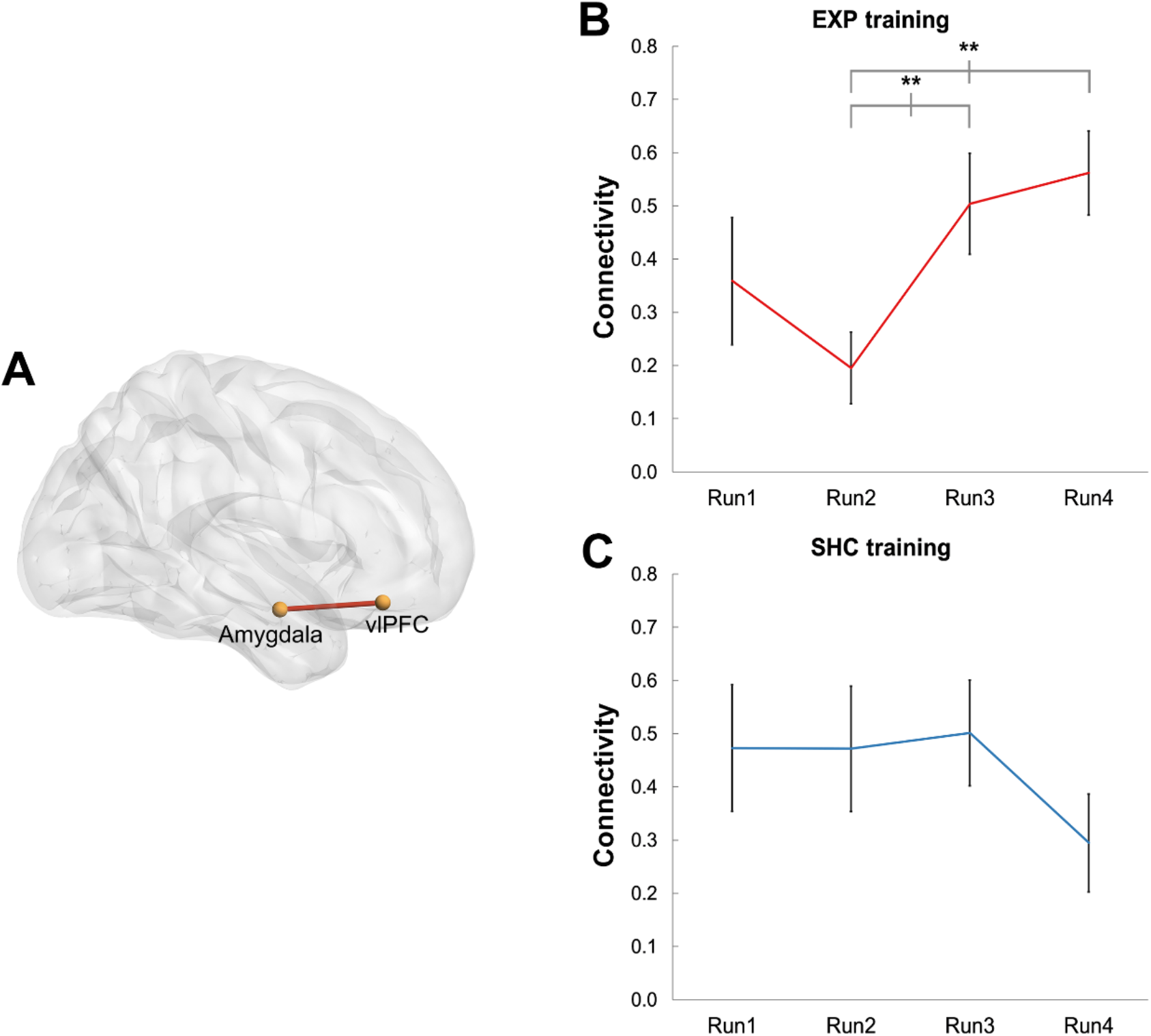
The emotion circuit trained in experimental training session (EXP, Panel A) which showed significant increased functional connectivity during emotion regulation after two runs of NFT training (Panel B). The controlled training (SHC) did not change the connectivity strength in this circuit (Panel C). Significant difference between training runs as tested by paired t-test are marked with asterisks (p < 0.01, two-tailed).

### Training-induced changes in anxiety

One outlier (> 2 SD from mean) for both, EXP and SHC sessions was excluded. Next, examining training-associated changes in the levels of anxious arousal, as assessed by VAS for EXP and SHC, revealed a significant decrease in VAS-rated anxiety levels following EXP training (Pre vs. Post training, t_20_ = 2.27, p = 0.035, Cohen’s *d* = 0.43, mean decrease 42.9% 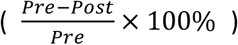 anxiety decrease, **Figure 4**), whereas for the sham training no significant changes were observed (t_21_ = −0.152, p = 0.881). Pre-training anxiety levels did not differ significantly between sessions (t_21_ = 0.60, p = 0.552).

**Figure 4.**
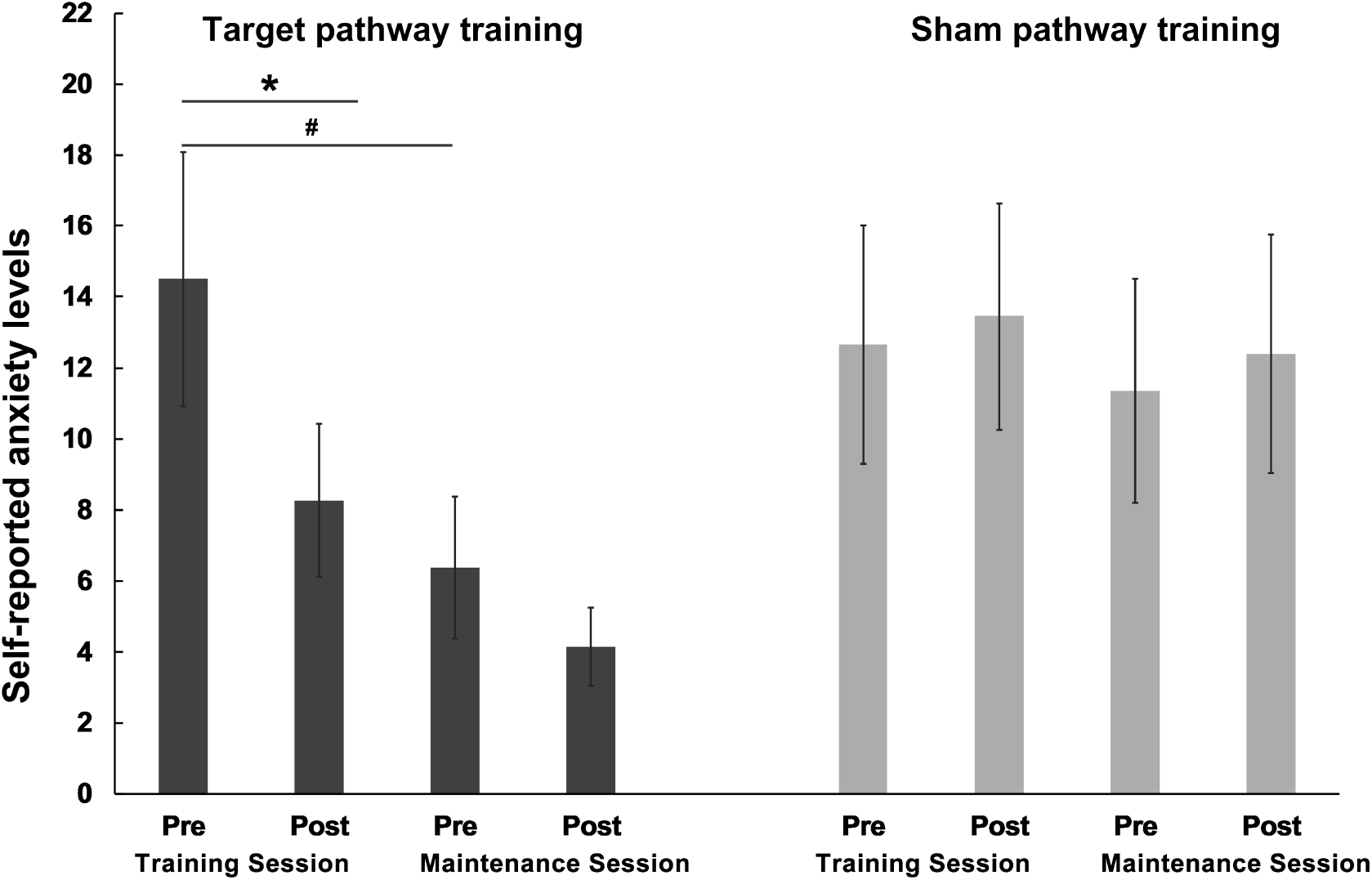
Self-reported anxiety level (VAS rating) decreased after EXP training (first two columns in dark gray) but not after SHC training (first two columns in light gray). Decreased anxiety level was maintained on Day 3 for EXP session (maintenance runs, last two columns in left and right panel). Differences in anxiety between pre and post training were tested by paired t-tests, two tailed. *p < 0.05. # denotes marginal significance, p < 0.10

### Association between neural and behavioral training success

A correlation analysis examined whether the anxiety level decrease after emotion regulation training related to the training-induced changes in the emotion regulation circuit. As in our previous study, differences between early and late training runs were considered as a neural index of individual training success (31). Results indicated that on an individual level the EXP training-associated anxiety decrease associated positively with the functional connectivity strength increase in the target pathway (r_20_ = 0.51, p = 0.016, Spearman).

### Transfer and maintenance of training success

Comparison of the mean amygdala-vlPFC connectivity between the successful training runs on Day 1 (Run3 and Run4) and the two transfer runs on Day 3 was used as an index of training maintenance effects (for a comparable strategy, see (31)). The analysis revealed no significant difference (p = 0.793) between the effective NFT-EXP runs and the M-EXP runs. Furthermore, there was a trend-significant correlation between the functional connectivity during NFT-EXP and M-EXP runs (r_21_ = 0.39, p = 0.068), suggesting that training success on the neural level in terms of regulatory control over the target pathway can be maintained independent of feedback and for a period of up-to three days.

After excluding two additional subjects based on their anxiety ratings on Day 3 (> 2 SD from mean), no significant training induced changes in anxiety levels were observed on Day 3 (Pre vs. Post for M-EXP, t_19_ = 1.04, p = 0.314; for M-SHC, t_22_ = −1.12, p = 0.274). However, plotting the anxiety ratings revealed attenuated anxiety levels at maintenance assessment following the EXP training session, but not the SHC (**Figure 4**). This pattern was further reflected in a marginal significant decrease in baseline anxiety levels on Day 1 versus Day 3 (Pre NFT-EXP vs. Pre M-EXP, t_19_ = 1.84, p = 0.082), while no changes were observed after the sham training session (Pre NFT-SHC vs. Pre M-SHC, t_22_ = 0.73, p = 0.475, **Figure 4**).

### Exploratory analysis: responders versus non-responders

We further explored the rate of non-responders (criteria: no improvements in primary neural and behavioral outcomes, n = 4) and evaluated the training success in the responders (n = 19), suggesting that determining putative responders may increase training success (details in **Supplementary Materials**).

## Discussion

The present proof-of-concept study employed a randomized, sham-controlled, within-subject design to evaluate the feasibility, functional relevance and maintenance of a novel connectivity-based rt-fMRI NF approach as a strategy to strengthen emotion regulation and decrease anxiety. During training of the amygdala-vlPFC pathway, but not the sham-control motor pathway, participants with high anxiety gained regulatory control over this ER-relevant pathway in terms of successful increasing functional connectivity strength over four subsequent training runs. On the behavioral level training of the target pathway – but not the sham pathway – was accompanied post-training by decreased anxious arousal ratings. On the individual level, the neural and behavioral indices of training success were significantly positively associated, further confirming the functional relevance of successful amygdala-vlPFC connectivity regulation. Finally, training success in terms of regulatory control over the amygdala-vlPFC pathway was maintained in the absence of feedback and for a period of three days, with preliminary evidence suggesting that the anxiety reduction was partly maintained. Importantly, no changes in the primary neural and behavioral outcomes were observed during the sham condition, arguing against unspecific effects of training or simple habituation.

The target amygdala-vlPFC pathway in the present study has previously been demonstrated to play an important role in successful ER (5, 15, 16) with rt-fMRI NF studies suggesting that successful down-regulation over regional amygdala activity associates with both, increased connectivity in the amygdala-prefrontal pathways as well as enhanced ER (38, 39). Moreover, previous clinical studies have emphasized the relevance of the amygdala-PFC circuits for treatment success, with changes in this pathway predictive of symptom-reduction after cognitive behavioral therapy (CBT) (20) and anxiolytic drug treatment (21, 22) in patients with exaggerated anxiety. In line with our hypothesis, successful training of the target pathway resulted in associated-decreases in anxiety ratings thereby confirming both the important role of the amygdala-vlPFC pathway in the regulation of anxiety as well as the functional relevance of the training. To increase the clinical relevance of the present proof-of-concept study subjects with high anxiety were recruited, and the training-associated decrease in anxious-arousal thus suggests that amygdala-vlPFC training may have to potential to normalize deficient prefrontal control of the amygdala and exaggerated levels of anxiety in clinical populations.

Of particular relevance for the application of NF-training approaches in clinical practice (30, 60, 61), the present study observed that subjects were able to maintain the control over the emotion regulation pathway and its effect on anxiety decrease in the absence of feedback and for a period of at least three days. These findings are in line with previous studies evaluating transfer and maintenance effects of rt-fMRI NF-assisted control over regional brain activity (31, 62), and additionally suggest that successful neuro-modulatory control on the pathway-level can last beyond the duration of the initial training and thus transfer to contexts outside of the MRI-environment.

Despite increasing interest in the application of rt-fMRI only a few studies to date have directly evaluated effects of neurofeedback training on functional connectivity between brain regions (50, 63–68). In line with the present findings, a previous study with a relatively small sample of healthy subjects revealed initial evidence for the feasibility of connectivity-informed NF which was associated with increased perception of positive valence stimuli. Importantly, the present study demonstrated the efficacy of this approach in decreasing anxious arousal, a transdiagnostic psychopathology marker (3), in subjects with high anxiety levels and thus may represent and important initial step towards the clinical application of connectivity-informed NF.

While these initially promising evaluations of functional connectivity-based NF training approaches are encouraging, there is still considerable room for improvement to promote transfer into clinical practice. For instance, the present study found preliminary evidence that anxiety attenuation was maintained after three days, however no further reduction was observed during the transfer session. Future studies should explore whether improved training strategies (52) and more intense or longer training schedules may lead to more robust and enduring behavioral effects in the absence of neurofeedback. Moreover, a considerable inter-individual variance in the neural and behavioral indices of training success were observed suggesting that some individuals are more likely to benefit from functional - connectivity-based training than others. Future studies are needed to identify optimal neural or behavioral predictors of training success allowing better selection of individuals who may benefit most from rt-fMRI NF training approaches. Recent findings suggest that baseline anxiety (66) or behavioral performance (69) may represent promising behavioral markers, although robust training-success predictors on the neural level remain to be determined.

Although the within-subject design, inclusion of a sham-control training and the preregistration of the primary outcomes permitted a rigorous control for a number of potential confounds, the present findings still need to be considered in the context of some limitations. First, to allow an evaluation of the training independent of menstrual-cycle or gender effects on ER and associated neural activity (46–48), our proof-of-concept study focused on male participants. The question of whether training success generalizes to female subjects and potential sex-differences therefore remains to be addressed. For clinical practice it will be important to compare both training and functional outcome success using activity- and connectivity-based feedback approaches. Connectivity-based NF comes at the cost of longer delay times and higher dimensionality (68) such that learning with this signal may be more demanding, possibly limiting efficacy in patients with cognitive impairments.

In summary, the present findings demonstrate that real-time functional connectivity-based neurofeedback training is feasible and targeting amygdala-prefrontal pathways with this training may represent a potential strategy to decrease anxiety in clinical populations. Importantly, neural and behavioral training success was maintained in the absence of feedback-guidance and for a period of at least three days.

## Funding and disclosure

This work was supported by the National Natural Science Foundation of China (NSFC, 91632117 to BB; 31530032 to KK), Fundamental Research Funds for the Central Universities of China (ZYGX2015Z002 to BB) and the Sichuan Science and Technology Department (2018JY0001 to BB).

The authors report no conflicts of interest.

